# Multimodal mechano-microscopy reveals mechanical phenotypes of breast cancer spheroids in three dimensions

**DOI:** 10.1101/2024.04.05.588260

**Authors:** Alireza Mowla, Matt S. Hepburn, Jiayue Li, Danielle Vahala, Sebastian E. Amos, Liisa M. Hirvonen, Rowan W. Sanderson, Philip Wijesinghe, Samuel Maher, Yu Suk Choi, Brendan F. Kennedy

## Abstract

Cancer cell invasion relies on an equilibrium between cell deformability and the biophysical constraints imposed by the extracellular matrix (ECM). However, there is little consensus on the nature of the local biomechanical alterations in cancer cell dissemination in the context of three-dimensional (3D) tumor microenvironments (TME). While the shortcomings of two-dimensional (2D) models in replicating *in situ* cell behavior are well known, 3D TME models remain underutilized because contemporary mechanical quantification tools are limited to surface measurements. Here, we overcome this major challenge by quantifying local mechanics of cancer cell spheroids in 3D TMEs. We achieve this using multimodal mechano-microscopy, integrating optical coherence microscopy-based elasticity imaging with confocal fluorescence microscopy. We observe that non-metastatic cancer spheroids show no invasion while showing increased peripheral cell elasticity in both stiff and soft environments. Metastatic cancer spheroids, however, show ECM-mediated softening in a stiff microenvironment and, in a soft environment, initiate cell invasion with peripheral softening associated with early metastatic dissemination. This exemplar of live-cell 3D mechanotyping supports that invasion increases cell deformability in a 3D context, illustrating the power of multimodal mechano-microscopy for quantitative mechanobiology *in situ*.

## I. INTRODUCTION

During metastasis, the main cause of cancer-related deaths, cancer cells must overcome mechanical challenges to invade distant tissues and organs [1]. Understanding the physiochemical interactions between the tumor microenvironment (TME) and cancer cells is imperative in developing novel anticancer treatments [2]. Although it is known that various cancer types influence metastasis through a range of mechanotransduction pathways [3], the lack of tools to measure subcellular elasticity (*i*.*e*., Young’s modulus) in three-dimensional (3D) *in vitro* tumor models, such as widely used cell spheroid systems, impedes current attempts to link functional biochemistry to the local biomechanical phenotypes of cancer [4]. Atomic force microscopy (AFM), a gold standard in mechanobiology, does not offer depth penetration [5]. Micropipette aspiration, parallel-plate compression [5] and deformability cytometry [6] provide bulk mechanical measurements of entire cells, and optical tweezers and stress mechano-sensors provide invasive point measurements [5]. 3D traction force microscopy has shown great promise, but provides measurements of forces exerted by cells, rather than a mechanical property, such as Young’s modulus, of cells and their surrounding environment [7]. Optical elastography techniques, particularly optical coherence elastography (OCE) and Brillouin microscopy, have shown promise in biomechanical characterization of cells, biomaterials, and tissues, as well as advancement towards clinical applications [8–10]. However, optical elastography techniques have key limitations. For example, OCE, which is based on optical coherence tomography (OCT), is unable to achieve subcellular elasticity resolution in both individual cells [11–13] and cell spheroid models [14]. Brillouin microscopy can achieve subcellular resolution, however, it measures longitudinal modulus in the GHz regime, which is challenging to relate to Young’s modulus, the most commonly used mechanical modulus, independent of water content and, also, has a limited penetration depth (∼100–200 μm) in opaque samples [15]. Furthermore, as it is becoming increasingly clear that the interplay between structure, biomechanics and biochemistry is central to the onset and progression of cancer [16], it is likely that a multimodal imaging platform is required to enable future breakthroughs in mechanobiology, yet no such platform currently exists.

The paucity of tools available to non-invasively map 3D Young’s modulus of TMEs leaves the mechanical phenotype of metastasis and the role of extracellular matrix (ECM) elasticity subject to debate [5]. Although metastatic cells are usually reported to be softer than their non-malignant counterparts [17], studies also suggest that they can instead stiffen during invasion [18]. Furthermore, while there is a positive correlation between increased ECM elasticity and cell invasion [19], it has also been reported that a stiff ECM indirectly inhibits cancer cell metastasis by modulation of mesenchymal stem cell secretions following mechanically induced differentiation [20]. These studies typically rely on point or 2D mechanical characterization of isolated cell systems that are well known to exhibit behaviors atypical of *in vivo* settings. Here, we overcome this major limitation by developing multimodal mechano-microscopy, an imaging platform that is capable of non-invasively quantifying Young’s modulus with subcellular resolution, co-registered with subcellular function provided by integrated confocal fluorescence microscopy (CFM), as well as 3D micro-scale structure from optical coherence microscopy (OCM). Mechano-microscopy is a low-coherence interferometric technique that provides a penetration depth of ∼500 μm in scattering samples [21]. Mechano-microscopy uses phase-sensitive OCM to map nano- to micro-scale axial deformation induced in the sample using a piezoelectric actuator with spatial resolution of 0.5×0.5×1.4 μm^3^ (*xyz*) in air. In post-processing, a mechanical model based on continuum mechanics is used to convert experimentally measured deformation into a high-resolution map of 3D Young’s modulus. Here, we demonstrate the capability of multimodal mechano-microscopy by performing a study on non-metastatic and metastatic cancer cell spheroids. Through close correspondence between structural OCM, Young’s modulus and CFM images, our results confirm that the mechanical phenotype of metastasizing cancer cells encapsulated in stiff and soft 3D TME-mimicking hydrogels, representative of primary and invasive sites [22], respectively, is mediated by the ECM stiffness. We believe that the new capability of multimodal mechano-microscopy, demonstrated here, is primed for the next generation of discoveries in cancer mechanobiology.

## II. RESULTS

Mechano-microscopy quantifies the local micro-scale mechanical properties of a sample using the principles of compression OCE [23,24]. As illustrated in the schematic in Fig. 1(a), the sample is placed on a microscope coverslip. The side distal to the incident optical beam is contacted by a compliant silicone layer [25], followed by a piezoelectric actuator that imparts sequential micro-scale compression to the system. The total deformation of the sample is less than 0.1% of its original thickness. The deformation is imaged using OCM with spatial resolution of 0.5×0.5×1.4 μm^3^ (*xyz*) in air. The local Young’s modulus is estimated at a system resolution of 5×5×15 μm^3^ (*xyz*) from deformation maps with reference to the estimated stress of the compliant silicone layer (details are presented in Section IV. A). Multimodal mechano-microscopy further records co-localized sample structure from the OCM signal-to-noise ratio (SNR) and fluorescence from CFM. We first validate our measurements against AFM on blank, homogeneous gelatin methacryloyl (GelMA) samples (*n*=2) that were photopolymerized for 15 and 60 seconds, respectively, hereby referred to as soft and stiff hydrogels. Figure 1(b) shows that the Young’s modulus of the stiff hydrogels is ∼3.5× greater than that of the soft hydrogels, with close correspondence between mechano-microscopy and AFM. Mechano-microscopy measures a slightly lower mean Young’s modulus of both soft and stiff hydrogels than AFM. Because of the different mechanical loading, such inter-tool variability is expected and well-documented for biological samples on this scale [26]. Figures 1(c) and 1(d) exemplify *xy* (*en face*) and *yz* cross-sectional images of local Young’s modulus of a living breast cancer (MCF7) spheroid encapsulated in stiff GelMA. Figures 1(e)–1(g) present the full complement of imaging capabilities with 3D visualizations of structure, Young’s modulus and fluorescence over a volume of 300×300×160 μm^3^ (*xyz*). The cyan and red regions in Fig. 1(g) represent live-stained nuclei (Hoechst 33342) and cell membranes (CellMask Green), respectively (details are presented in Section IV. F).

**Fig. 1.**
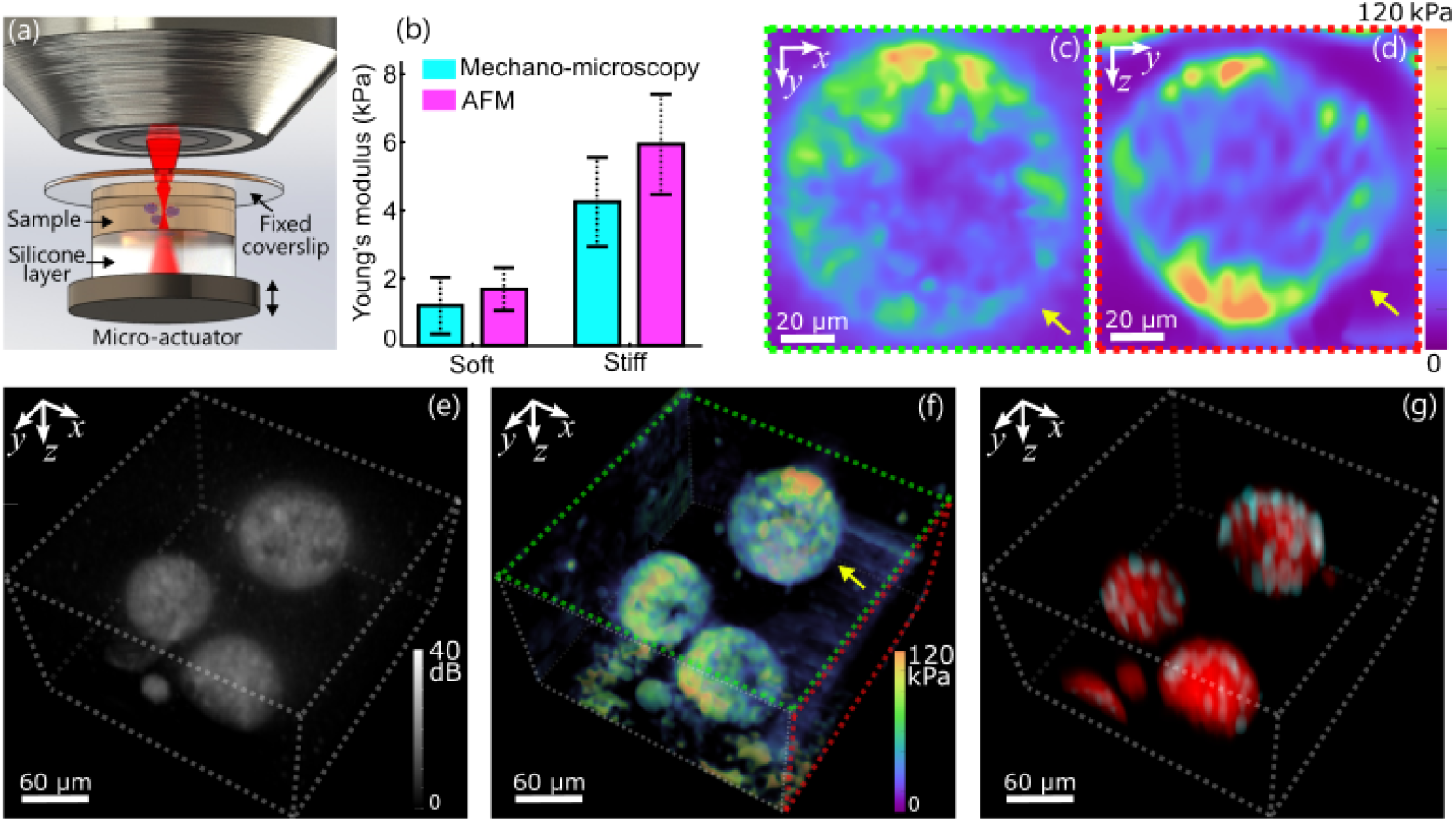
Multimodal mechano-microscopy. (a) Schematic of the sample setup. (b) Young’s modulus of soft and stiff blank GelMA samples (*n*=2) measured using mechano-microscopy (volumetric measurements) and AFM (surface measurements). Error bars represent standard deviation across the sampled field-of-view. (c) *En face* and (d) B-scan Young’s modulus of a non-metastatic cell spheroid. (e–g) Volumetric maps of non-metastatic breast cancer cell spheroids acquired by multimodal mechano-microscopy, where (e) OCM SNR, (f) Young’s modulus and (g) fluorescence labelled with nuclear (cyan) and membrane (red) fluorescent dyes. Yellow arrows indicate the same spheroid in (c), (d) and (f).

In Fig. 2, a comparison is presented between compression OCE (details are presented in Section IV. C) [11] and multimodal mechano-microscopy performed on breast cancer (MCF7) cell spheroids encapsulated in GelMA, similar to the spheroid presented in Figs. 1(c)–1(g). In Fig. 2(b), a magnified OCT image is presented that corresponds to the region highlighted by the cyan box in Fig. 2(a). From the co-registered fluorescence image presented in Fig. 2(f), it is evident that the higher resolution provided by OCM [Fig. 2(c)] reveals structures not visible in the OCT image. In particular, the cyan arrows in Figs. 2(c) and 2(f) highlight subcellular structures, likely corresponding to the cell nuclei, which present as regions of lower OCM SNR, due to relatively lower and less heterogenous refractive index and mass density than cytoplasm [27–29]. It is challenging to identify these structures in the OCT image in Fig. 2(b). The Young’s modulus images, corresponding to OCE and mechano-microscopy, respectively, are presented in Figs. 2(d) and 2(e), respectively. It is evident that, for these spheroids, it is not possible to reveal intra-spheroidal structures using OCE, with a relatively uniform Young’s modulus presented throughout the spheroids. Mechano-microscopy [Fig. 2(e)], on the other hand, reveals rich contrast within the spheroids. In particular, a relatively high Young’s modulus is evident around the perimeter of the spheroids, which, through comparison with the CFM image [Fig. 2(f)] likely corresponds to regions with a denser concentration of cell nuclei, which is typically the stiffest organelle within a cell [30].

**Fig. 2.**
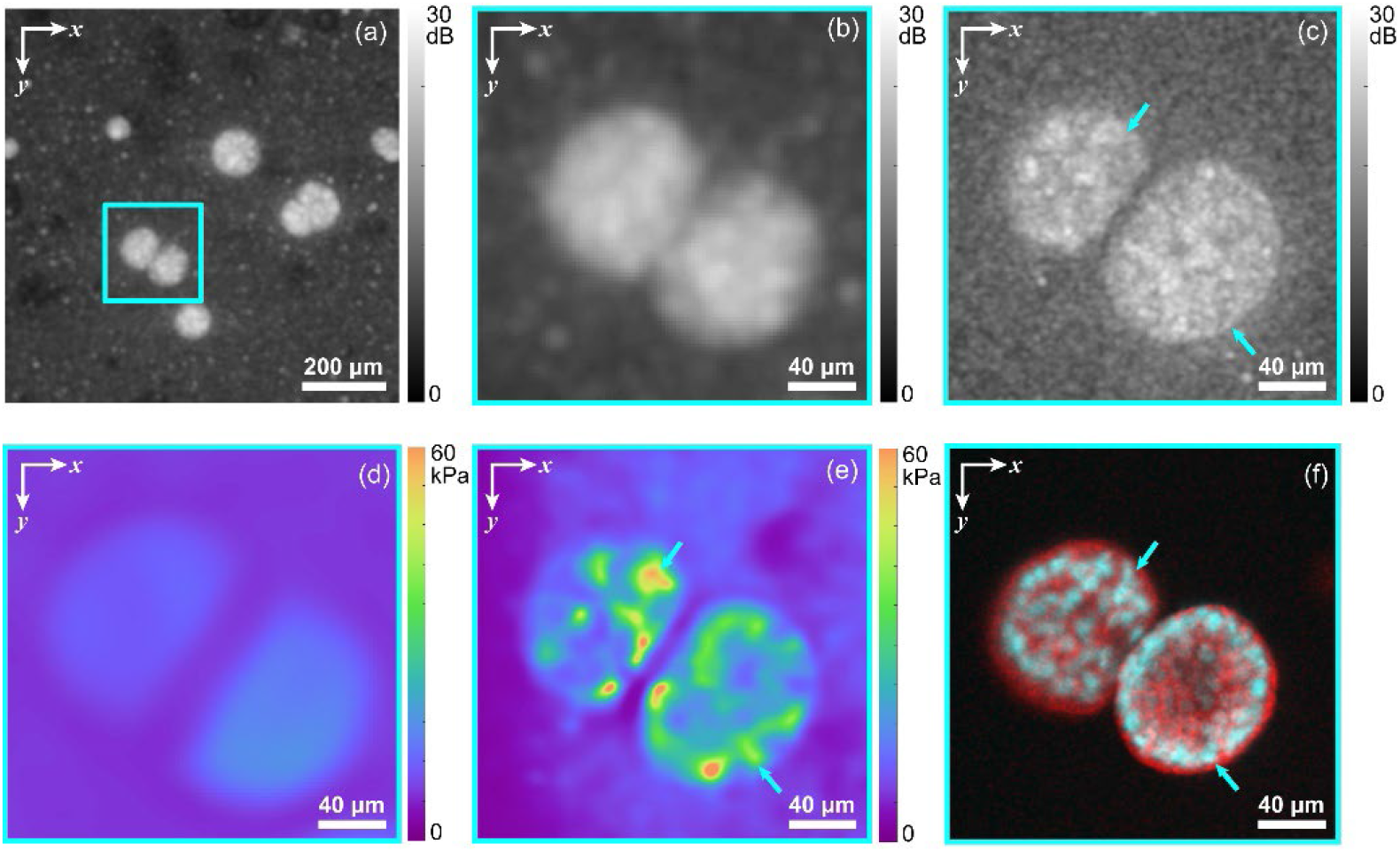
OCE vs mechano-microscopy of non-metastatic breast cancer cell spheroids in GelMA. (a) *En face* (*xy*) OCT image of the spheroids. The cyan box indicates the location of spheroids of interest. The corresponding magnified (b) OCT SNR and (d) Young’s modulus using OCE in comparison to the co-registered (c) OCM SNR, (e) Young’s modulus and (f) fluorescence labelled with nuclear (cyan) and membrane (red) fluorescent dyes using mechano-microscopy. Cyan arrows in (c), (e) and (f) highlight nuclei in the cell spheroids.

We demonstrate mechano-microscopy by investigating the mechanical phenotypes of non-metastatic (MCF7) and metastatic (MDA-MB-231) cancer cell spheroids cultured for 6–8 days in stiff (∼5 kPa) and soft (∼1.5 kPa) 3D GelMA hydrogels. In Fig. 3, examples of non-metastatic spheroids cultured in both the stiff and soft hydrogels are presented. *En face* (*xy)* cross-sections of OCM [Figs. 3(a) and 3(d)], Young’s modulus [Figs. 3(b) and 3(e)] and fluorescence [Figs. 3(c) and 3(f)] are presented at three depths (the middle depth represents the spheroid’s central cross-section), representative of repeated independent samples (*n*=4). In the fluorescence image, the expected initiation of hypoxic conditions and necrotic-like phenotypes in the core (cyan arrows) suggests greater cell concentration in the spheroid periphery (magenta arrows) [31]. This has been observed previously at lower resolution using OCE [14] and may be associated with the formation of growth-arrested clusters, which are similar to acini found *in vivo* in mammary glands [32]. In the OCM images, as in Figs. 2(a) and 2(d), nuclei present as structures with low SNR, suggesting that they exhibit lower optical backscattering at the wavelengths used than the higher SNR observed in the cell body. In the Young’s modulus images [Figs. 3(b) and 3(e)], elevated elasticity is again observed at the spheroid periphery. Cancer cells are characterized by increased deposition and remodeling of ECM proteins, leading to potential self-compression of the tumor and the observed peripheral stiffening [1]. The spheroid encapsulated in the soft hydrogel [Figs. 3(d)–3(f)] is larger than the one in the stiff hydrogel [Figs. 3(a)–3(c)], consistent with our previous study using OCE, which showed that spheroids grown in soft hydrogels have relatively larger volumes than those grown in stiff hydrogels [14].

**Fig. 3.**
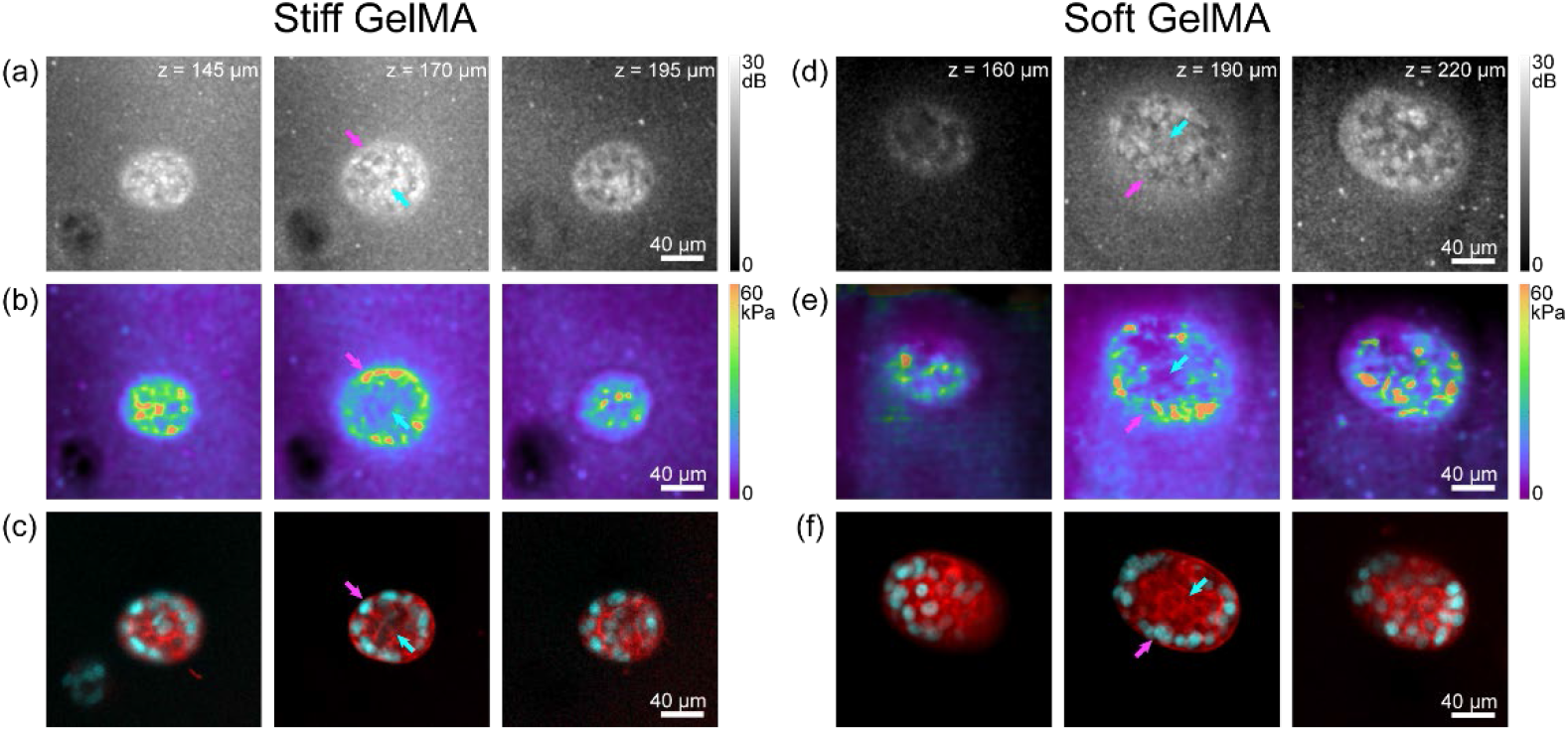
Multimodal mechano-microscopy of non-metastatic breast cancer cell spheroids in stiff and soft GelMA. *En face* images of (a, d) OCM SNR, (b, e) Young’s modulus and the co-registered (c, f) fluorescence labelled with nuclear (cyan) and membrane (red) fluorescent dyes, presented at three annotated depths, corresponding to the top, middle and bottom cross-sectional planes of the cell spheroids. Cyan and magenta arrows in (a)–(f) indicate the core and periphery of the cell spheroids, respectively.

In Fig. 4, examples of metastatic spheroids cultured in both the stiff [Figs. 4(a)–4(c)] and soft [Figs. 4(d)–4(f)] hydrogels are presented. Interestingly, metastatic spheroids in a stiff environment do not present cell dissemination. Only when such spheroids are in a softer environment can early metastatic dissemination be observed. Metastatic spheroids in stiff hydrogels present a similar spatial distribution of Young’s modulus to that of the non-metastatic spheroids; however, they demonstrate an overall lower Young’s modulus [Fig. 4(b)]. In addition, the peripheral cell membrane and ECM border exhibit disorganization, and an overall higher density of nuclei in the core [Figs. 4(a)–4(c)], compared to their non-metastatic counterparts. Distinct to all the other cases, dissemination of metastatic spheroids in a soft environment is contrasted by a softening of the peripheral cells compared to the core and an associated decrease in peripheral nuclear density [Figs. 4(d)–4(f)], possibly induced by increased nuclear volume in disseminating cells [33]. Although cell invasion has been suggested in stiffer and more constricting ECM environments, the associated mechanisms are still poorly understood in 3D contexts, emphasizing the present need to disentangle the roles of ECM pore size, density and local Young’s modulus from the mechano-phenotype [1,5]. Interestingly, a similar reduction in invasive phenotypes in stiff 3D hydrogels has been observed in 2D images of Brillouin frequency shift and, also, with implantable sensors [34], giving promise to 3D models unifying our understanding of cancer invasion.

**Fig. 4.**
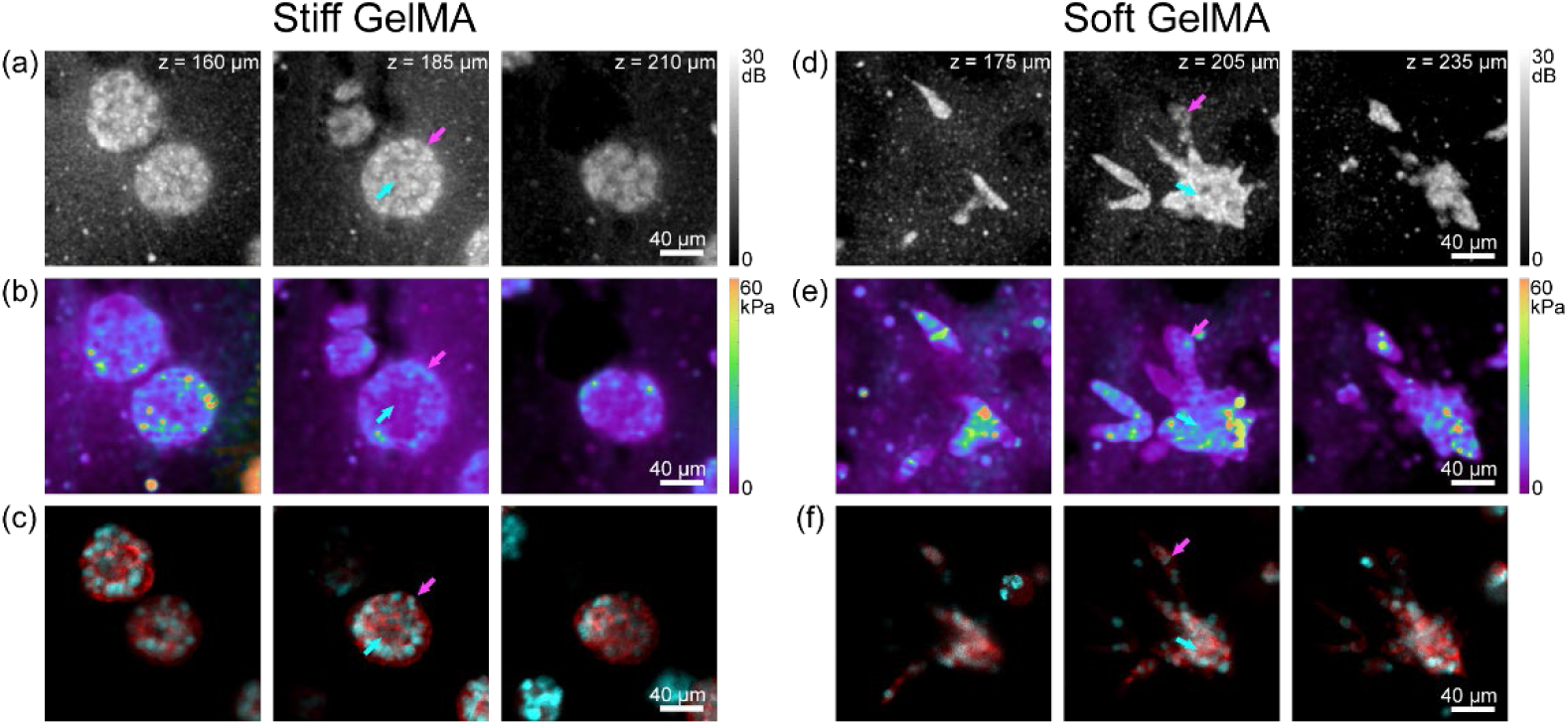
Multimodal mechano-microscopy of metastatic breast cancer cell spheroids in stiff and soft GelMA. *En face* images of (a, d) OCM SNR, (b, e) Young’s modulus and the co-registered (c, f) fluorescence labelled with nuclear (cyan) and membrane (red) fluorescent dyes, presented at three annotated depths, corresponding to the top, middle and bottom cross-sectional planes of the cell spheroids. Cyan and magenta arrows in (a)–(f) indicate the core and periphery of the cell spheroids, respectively.

The Young’s moduli in the periphery and core were quantified using automated segmentation in 3D (details are presented in Section IV. G) for all samples (*n*=4 or 5) [Fig. 5(a)], representing a statistical validation of our observations. Furthermore, the nuclei density in the core and periphery regions was quantified by their mean normalized fluorescent intensity, as presented in Fig. 5(b) (details are presented in Section IV. G), showing an increased cell density at the periphery in non-metastatic spheroids while indicating reduced cell density at the core in metastatic spheroids embedded in soft GelMA.

**Fig. 5.**
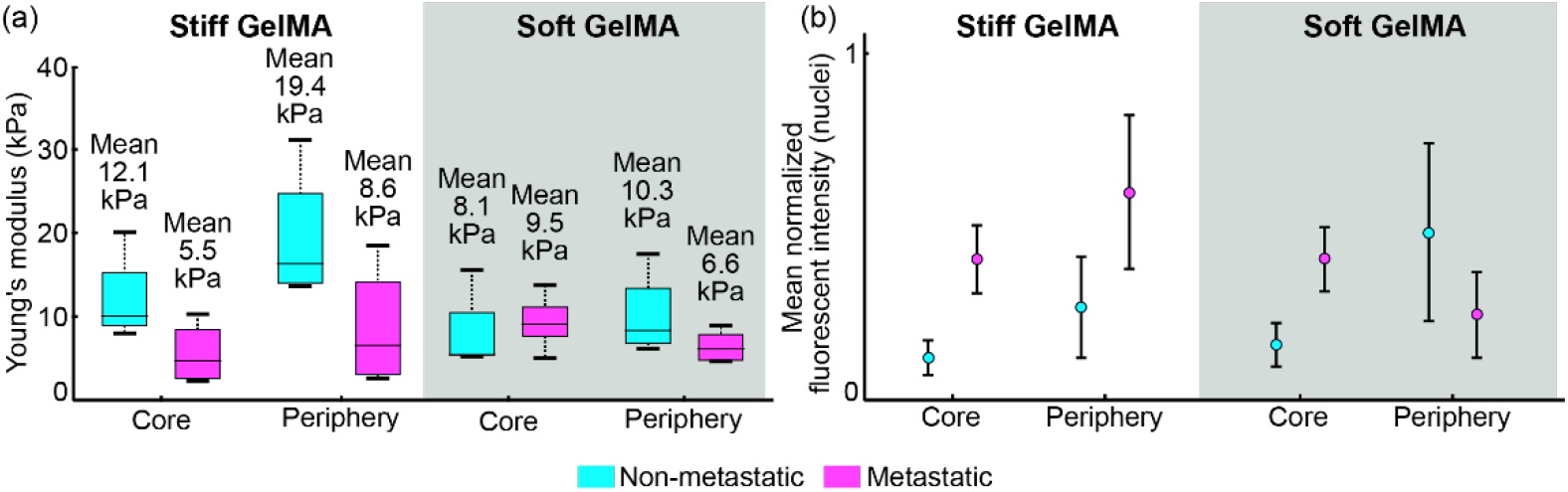
(a) Young’s modulus distribution at the core and periphery regions in all samples (*n*=4 or 5). (b) The nuclei density in the corresponding core/periphery in Figs. 3(c) and 3(f), and Figs. 4(c) and 4(f), respectively, quantified by the mean normalized fluorescent intensity of the nucleus.

## III. DISCUSSION

In this paper, we have introduced multimodal mechano-microscopy, a unique imaging platform that provides micro-scale, 3D images of the structure, biomechanics and biochemistry of living tumor spheroids encapsulated in GelMA hydrogels. In this initial demonstration, multimodal mechano-microscopy was used to provide insight in the development of non-metastatic and metastatic tumor spheroids in both stiff and soft environments. Importantly, through direct comparison between metastatic tumor spheroids in stiff and soft GelMA, we have demonstrated that cancer invasion highlighted by the dissemination of metastatic spheroids is regulated by the ECM stiffness and is facilitated by softening of the matrix and metastatic cancer cells in the periphery. Given the importance of reconciling these properties in most *in vitro* models, we believe that our imaging platform will find broad application in other areas, including in imaging other types of spheroids, as well as single cells embedded in biomaterials and tissue organoids. Additionally, analogously to related OCE techniques, multimodal mechano-microscopy can be used to provide insight in tissue mechanics, with higher spatial resolution in 3D than is currently possible.

In this study, we have demonstrated that CFM, integrated with mechano-microscopy, enables co-registered OCM, mechano-microscopy and fluorescence images to relate subcellular structures with localized regions of Young’s modulus within spheroids in 3D. CFM was performed directly after the acquisition of OCM/mechano-microscopy data. This time delay between the different imaging modes, although relatively small, may cause slight discrepancies in image co-registration. It would be straightforward to address this in future work to achieve temporal overlap in the acquisition of each image in multimodal mechano-microscopy. This would facilitate the imaging of spheroids at different time points in their development, which could provide important insight in tumor development. An additional limitation for time-lapsed imaging in the current setup is the lack of control of environmental factors, such as temperature, humidity, CO_2_ and O_2_. To address this, the imaging platform could be developed towards high-throughput mechanical phenotyping and drug screening by integrating a customized stage-top incubator [35], combined with an automated wide-field scanning system [36] to accelerate the speed of volumetric imaging over the entire sample area. Staining cell nuclei and membranes in live spheroids effectively visualizes intra-spheroid morphological variation and subcellular functional deformation associated with mechanical phenotypes of cancer spheroids. However, further insight could be gained by staining other important organelles associated with mechanotransduction, such as actin filaments and focal adhesions, that drive the mechanobiology of cancer cells [37,38]. In addition, for long-term monitoring of spheroids in customized stage-top incubators, suggested above, one can transfect cells with fluorescent proteins. This would provide stable and long-lasting fluorescent markers integrated within the cells’ natural protein structures [39]. Furthermore, incorporating extra excitation wavelengths, such as 561 nm, in addition to 405 nm and 488 nm channels, would provide greater flexibility for using a wider range of fluorescent probes.

The central wavelength of the light source used in our system is ∼800 nm, providing a measured OCM axial resolution of 1.4 μm in air. It has been reported that visible-light OCM, where the central wavelength is in the visible portion of the electromagnetic spectrum, can provide sub-micrometer axial resolution [40]. However, a disadvantage of this approach is the high optical attenuation at these lower wavelengths, which limits the penetration depth in turbid samples. For this reason, we choose a longer wavelength, trading off axial resolution for increased penetration depth, enabling us to image through entire tumor spheroids.

To estimate Young’s modulus of the sample, the mechanical model used in mechano-microscopy, as is the case for most elastography techniques, requires simplifying assumptions on the nature of the sample’s properties and its deformation. In this study, to minimize the hyperelastic effect of the sample on the measurements of Young’s modulus [41], a minimum pre-strain (<5%), *i*.*e*., a bulk strain applied prior to the micro-scale actuation, was used to ensure uniform contact between the sample, layer and compression plates. Furthermore, to reduce viscoelastic effects, the imaging protocol was carefully tailored by encapsulating cell spheroids into GelMA that exhibits low viscoelasticity [42], and the samples were deformed at a quasi-static loading frequency [11] to ensure the instantaneous elastic strain was measured. However, a more generalized mechanical model could be developed to account for viscoelasticity [43,44] of the sample to enable multiparametric quantitative characterization of spheroid mechanics.

In summary, multimodal mechano-microscopy presents a new opportunity for quantifying local Young’s modulus in live 3D samples with subcellular resolution. In contrast to previous methods, we can directly reveal mechanical phenotypes *in situ*, which is a major step towards disambiguating physiologically relevant modifications in cell mechanics from those induced by point contact or by atypical 2D culture. Beyond this example, mechano-microscopy is ready to explore a host of cancer types and TME models in 3D and is well-poised to propel mechanobiology towards the quantification and specificity needed to understand complex biophysical processes in disease and development.

## IV. METHODS

### A. Multimodal mechano-microscopy

Multimodal mechano-microscopy integrates high-resolution interferometric detection of OCM, a high-resolution variant of OCT [21], with compression OCE [25] and CFM. A schematic diagram of the imaging system is shown in Fig. 6. The system uses a supercontinuum laser (SuperK Extreme EXW-4 OCT, NKT Photonics, Denmark), whose output was shaped to a spectral range of 650 nm to 950 nm using a combination of a long-pass dichroic mirror (DMLP950, Thorlabs Inc., USA), a short-pass dichroic mirror (DMSP650, Thorlabs Inc., USA), a long-pass filter (FELH0650, Thorlabs Inc., USA) and a short-pass filter (FESH0950, Thorlabs Inc., USA). This spectrum corresponded to a full-width at half-maximum (FWHM) bandwidth of ∼250 nm, providing a measured OCM axial resolution of 1.4 μm in air. The system was implemented as a Michelson interferometer in a dual-arm configuration with the same optics in the reference beam path to match optical dispersion. A dispersion compensating block was used to account for residual dispersion. The sample arm beam was expanded to fill the entrance pupil of the objective lens (20× 0.75 NA, CFI Plan Apo, Nikon) providing a measured lateral resolution of 0.5 μm. Scanning was achieved using a 2D galvanometer system (GVSM002-EC/M, Thorlabs Inc., USA).

**Fig. 6.**
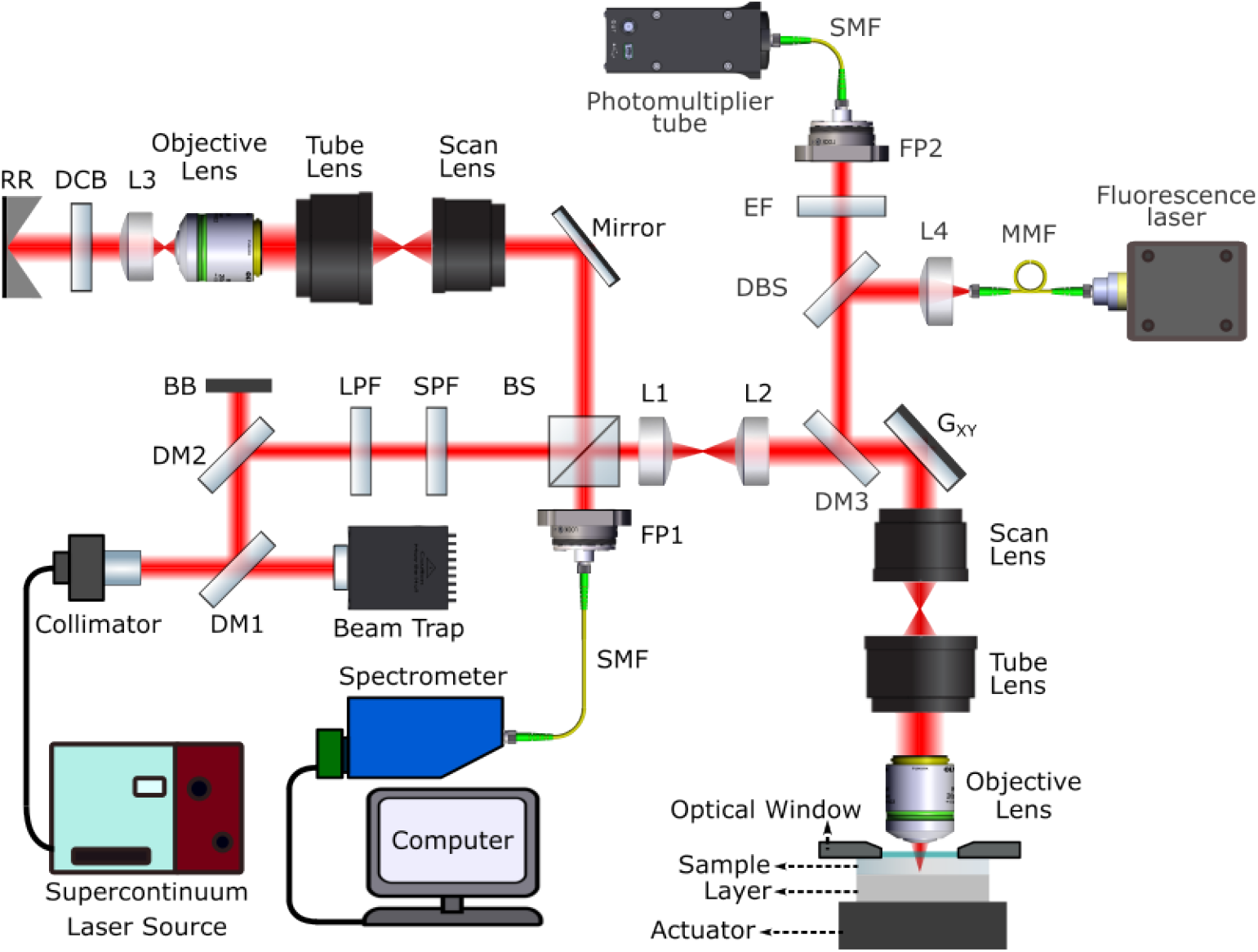
Schematic diagram of the multimodal mechano-microscopy system. RR: reference reflector, DCB: dispersion compensation block, L: lens, BS: beam splitter, SPF: short-pass filter, LPF: long-pass filter, BB: beam block, DM: dichroic mirror, FP: fiber port, SMF: single-mode optical fiber, G_xy_: *xy* galvanometer mirrors, DBS: dichroic beam-splitter, MMF: multi-mode optical fiber, EF: edge filter.

A sample with a thickness of ∼400–500 μm and a 1 mm thick compliant silicone layer were compressed between the coverslip and a rigid optical window attached to an annular piezoelectric actuator (Piezomechanik GmbH, Germany). We used a spectrometer comprising a 2048-pixel line camera with a maximum line rate of 130 kHz (Wasatch Photonics, USA) to detect the spectral interference at each *xy* location. The B-scan and volume acquisition times were 20.4 ms and 40.8 s, respectively. The annular actuator was driven in a quasi-static regime with a 24.5 Hz square wave, synchronized with the acquisition of B-scans. Two B-scans were acquired at each *y*-location such that one B-scan was acquired at the unloaded and one at the loaded state. Local displacement was calculated from the phase difference between the loaded and unloaded B-scans and strain was estimated as the gradient of the axial displacement with depth using weighted least squares linear regression [25]. Strain in the layer was related to stress through knowledge of the stress–strain response of the compliant layer in contact with the sample. By assuming uniaxial stress, the local stress at the surface of the sample was divided by the local strain in the sample to estimate sample elasticity as the tangent modulus, which is equivalent to Young’s modulus under the assumption of linear elasticity. An elasticity system resolution of 5×5×15 μm^3^ (*xyz*) is calculated as the convolution of the OCM system resolution and the FWHM of the signal processing used, *i*.*e*., Gaussian smoothing and weighted least squares linear regression [45]. To minimize the effect of friction on the precision [46] and accuracy [47] of the estimated Young’s modulus, customized hyaluronic acid (HA) solution was made to improve the lubrication of the contact surface between the window and GelMA hydrogels [48].

The integrated CFM system comprised two laser lines at 405 nm and 488 nm as the excitation light source (CNI Lasers, Changchun New Industries Optoelectronics Technology Co., China) and a photomultiplier tube (PMT2101/M, Thorlabs Inc., USA) as the detector. The CFM system used the same galvanometer scanning arm and optics as the OCM system to scan the focal point in the sample. The emitted fluorescence was focused into a single-mode optical fiber, which also acted as the confocal pinhole. The focal plane of the CFM system was aligned with the OCM system to co-register the focal planes of both imaging systems in the sample. The CFM point-spread function (PSF) was measured using small beads, and the resolution was determined from the FWHM of the PSF as 0.5 μm in the lateral direction and ∼8 μm in the axial direction. To obtain a *z*-stack and generate the 3D CFM image presented in Fig. 1(g), the sample mount setup was scanned in *z* in steps of 5 μm. The recorded fluorescence intensities at each scan point were saved into 64-bit, 2D binary images. Fiji software [49] was used to combine the two channels and multiple focal planes into 3D composite image stacks.

### B. Dynamic focusing to extend the depth of field

Mechano-microscopy uses a high numerical aperture (NA) microscope objective to achieve sub-micrometer lateral resolution. However, the high NA beam significantly decreases the depth of field, therefore, the imaging depth is limited to ∼10–15 μm around the focal plane. To overcome the limited depth of field and to extend the imaging depth, dynamic focusing was performed by scanning the sample at consecutive axial locations [50] and acquiring multiple volumetric scans. To generate the 3D visualizations in Figs. 1(e) and 1(f), 11 volumetric scans were acquired at partially overlapping focal depths along the *z*-axis inside the sample within ∼9 minutes. To vary the focus position in the sample, the sample mount assembly was translated in consecutive steps of 10 μm in air using a motorized translation stage. To generate the Young’s modulus image with extended depth of field as shown in Fig. 1(f), Young’s modulus was computed for each volumetric scan position, then, an axial Gaussian filter was applied with a peak corresponding to each volume’s focal plane. We summed the filtered Young’s modulus subvolumes to generate the extended depth of field 3D image. A similar approach was used to generate the 3D-OCM visualization in Fig. 1(e).

### C. Compression optical coherence elastography

To highlight the improved imaging capability of multimodal mechano-microscopy, as shown in Fig. 2, we compared its performance on cell spheroids with compression OCE based on a lower resolution OCT system demonstrated in a previous study on individual cells in 3D [11]. We pre-aligned the two systems and translated the sample setup from the multimodal mechano-microscopy to OCE, and performed consecutive scans of the same sample under the same loading conditions. The OCT system used is a fiber-based spectral-domain OCT system (Telesto 220, Thorlabs Inc., USA) that has a spectral bandwidth of 170 nm and a central wavelength of 1300 nm. The measured axial and lateral resolutions (FWHM) are 4.8 μm and 4.4 μm, respectively. Similar to mechano-microscopy, signal processing in OCE includes the combination of Gaussian smoothing and weighted least squares linear regression, resulting in an isotropic elasticity system resolution of 35 μm, calculated as the convolution of the OCT system resolution and the FWHM of the signal processing used.

### D. Hydrogel Fabrication and Cell Encapsulation

Lyophilised GelMA was initially placed in a desiccator vacuum with an attached filter cap (SteriFlip 0.22 μm, Millipore) to remove residual moisture. Following this, GelMA was weighed and dissolved in sterile Dulbecco’s Phosphate Buffered Saline (DPBS, Gibco) to make a 9% (w/v) GelMA solution. Additionally, a powdered photoinitator, Irgacure-2959 (2-Hydroxy-4’-(2-hydroxyethoxy)-2-methylpropiophenone, Sigma-Aldrich) dissolved in absolute ethanol was added to the solution at a final concentration of 0.1% (w/v). The solution was then protected from light and stored at 4°C overnight prior to use. On the day of encapsulation, the GelMA precursor solution was warmed to 37°C in a water bath for 1 hour. MCF-7 and MDA-MB-231 cells were resuspended in pre-warmed GelMA at 2000 cells/hydrogel. The cell-laden GelMA was then pipetted into circular 5 mm diameter, 300 μm thick polydimethylsiloxane (Wacker Chemie AG, Germany) molds atop a dichlorodimethylsilane-treated (Sigma-Aldrich) glass slide and covered with a methacrylated (3-(trimethoxysilyl) propylmethacrylate, Sigma-Aldrich) circular coverslip (10 mm diameter, Trajan). The cell-laden GelMA was then exposed to UV light (365 nm at ∼2.5 mw/cm^2^, Vilber ECX-F20.L) for 15 or 60 seconds, respectively, to produce soft or stiff gels. MCF7 and MDA-MB-231 spheroids were cultured for 16 (Fig. 3) and 6–8 (Fig. 4) days, respectively, to allow for adequate spheroid growth.

### E. Atomic force microscopy

An MFP-3D atomic force microscope (Asylum Research) equipped with 200 μm long gold-coated pyramidal-tipped pyrex-nitride cantilevers (PNP-TR, NanoWorld) was used to measure GelMA surface stiffness. Each sample was indented at five sites that were consistent between samples. Indentations were performed in triplicate at 2 nN while samples were immersed in PBS. Custom code in Igor Pro was used to analyze the linear portion of contact-generated force curves to derive Young’s Modulus. As presented in Fig. 1(b), AFM measured the Young’s Modulus of two soft GelMA hydrogels at 1.65±0.64 kPa and 1.73±0.61 kPa, and two stiff GelMA hydrogels at 5.23±2.28 kPa and 6.64±0.65 kPa, which are comparable to the volumetric Young’s Modulus measured by mechano-microscopy of two soft GelMA hydrogels at 1.34±0.93 kPa and 1.04±0.73 kPa, and two stiff GelMA hydrogels at 4.77±1.48 kPa and 3.72±1.12 kPa, respectively.

### F. Live cell staining protocol

The GelMA containing the cell spheroids, attached to coverslips, were kept on a 12-well plate in normal growth medium at 37°C 5% CO_2_ and stained just before imaging. CellMask Green (C37608, ThermoFisher) was mixed with 0.5 ml of growth medium in 1:200 dilution and Hoechst 33342 was added in 10 μM final concentration. The growth medium in the sample well was replaced with the dye mix, and the sample was incubated at 37°C for 20 minutes. The sample was then washed twice with warm PBS and returned to normal growth medium. The samples were imaged within 2 hours of staining.

### G. Statistical analysis

Young’s modulus in the core and periphery regions was determined by masking these regions in structural OCM volumes and then applying the masks to the corresponding Young’s modulus. The structural volumes were normalized to a range of 0 and 1 and then binarized with a threshold of 0.5. Regions that were disconnected from the main spheroid were removed and holes were filled to produce a single contiguous mask of each spheroid. Edge-smoothing was performed by convolving a 20×20-pixel (40×40 μm^2^) kernel of ones with the initial binary mask then thresholding out all values below 0.95 to create a smoothed binary mask. The centroid of each mask was determined by calculating the center of mass where each pixel in the binary mask was uniformly weighted. Each pixel was categorized according to its proximity to the centroid. For the non-aggressive spheroids in stiff GelMA, non-aggressive spheroids in soft GelMA, and aggressive spheroids in stiff GelMA, the core regions correspond to the 50% of pixels closest to the centroid, and the periphery regions correspond to the furthest 50% of pixels. Considering the approximate size of these spheroids, we chose the 50/50 volumetric portions for the core/periphery regions so that the outer cell layer with a radial length of ∼15 μm is covered in the outer 50% region. For the aggressive spheroids in soft GelMA, the core regions correspond to the nearest 75% of pixels, with the furthest 25% classified as periphery. In this case, we chose the 75/25 volumetric portions for the core/periphery regions so that the disseminating metastatic cell branches are covered in the outer 25% region. The mean Young’s modulus corresponding to the core and periphery regions of each spheroid was then determined by averaging each respective voxel in the core and periphery volumes. Finally, the averaged inner and peripheral Young’s modulus were determined for each individual spheroid.

The nuclei density in different spatial locations of the middle plane of the fluorescent image of the spheroids were quantified by measuring the mean fluorescent intensity of Hoechst 33342 from the spheroid center to the periphery region. This involved applying the same thresholding algorithm used in quantifying the elasticity distribution (described above). The mean intensity of the fluorescent nucleus channel was calculated by averaging the normalized cyan color value in the fluorescent image in each region.

## ACKNOWLEDGEMENTS

This study was supported by the Australian Research Council, The Ian Potter Foundation, Department of Health, Western Australia, Research Training Program Scholarship, Hackett Postgraduate Research Scholarship, and Cancer Council Western Australia. PW was supported by the 1851 Research Fellowship from the Royal Commission.

## AUTHOR DECLARATIONS

### Conflict of Interest

B.F.K. OncoRes Medical (F, I). The others have no conflicts to disclose.

### Ethics Approval

Ethics approval not required.

### Author Contributions

**Alireza Mowla:** Conceptualization (equal); Data curation (equal); Formal analysis (equal); Methodology (lead); Investigation (equal); Writing – original draft (lead); Writing – review & editing (equal). **Matt S. Hepburn:** Data curation (equal); Formal analysis (equal); Investigation (equal); Writing – review & editing (equal). **Jiayue Li:** Data curation (equal); Formal analysis (equal); Investigation (equal); Writing – review & editing (equal). **Danielle Vahala:** Data curation (equal); Formal analysis (supporting); Investigation (supporting); Writing – review & editing (equal). **Sebastian E. Amos:** Data curation (equal); Formal analysis (supporting); Investigation (supporting); Writing – review & editing (equal). **Liisa M. Hirvonen:** Data curation (supporting); Formal analysis (supporting); Investigation (supporting); Writing – review & editing (equal). **Rowan W. Sanderson:** Formal analysis (equal); Investigation (equal); Writing – review & editing (equal). **Philip Wijesinghe:** Formal analysis (equal); Investigation (equal); Writing – review & editing (equal). **Samuel Maher:** Data curation (equal); Formal analysis (supporting); Investigation (supporting); Writing – review & editing (equal). **Yu Suk Choi:** Conceptualization (equal); Funding acquisition (equal); Resources (equal); Writing – review & editing (equal). **Brendan F. Kennedy:** Conceptualization (equal); Funding acquisition (equal); Resources (equal); Writing – review & editing (equal).

## DATA AVAILABILITY

The data that support the findings of this study are available from the corresponding author upon reasonable request.

